# iDEP-assisted isolation of insulin secretory vesicles

**DOI:** 10.1101/2021.12.01.470798

**Authors:** Mahta Barekatain, Yameng Liu, Zhongying Wang, Vadim Cherezov, Scott E. Fraser, Kate L White, Mark A. Hayes

## Abstract

Organelle heterogeneity and inter-organelle associations within a single cell contribute to the limited sensitivity of current organelle separation techniques, thus hindering organelle subpopulation characterization. Here we use direct current insulator-based dielectrophoresis (DC-iDEP) as an unbiased separation method and demonstrate its capability by identifying distinct distribution patterns of insulin vesicles from pancreatic β-cells. A multiple voltage DC-iDEP strategy with increased range and sensitivity has been applied, and a differentiation factor (ratio of electrokinetic to dielectrophoretic mobility) has been used to characterize features of insulin vesicle distribution patterns. We observed a significant difference in the distribution pattern of insulin vesicles isolated from glucose-stimulated cells relative to unstimulated cells, in accordance with functional maturation of vesicles upon glucose stimulation, and interpret this to be indicative of high-resolution separation of vesicle subpopulation. DC-iDEP provides a path for future characterization of subtle biochemical differences of organelle subpopulations within any biological system.

## INTRODUCTION

Functional cell-to-cell heterogeneity within a given cell type arises from differences in genomic, epigenomic, transcriptomic, and proteomic^1^ components and their specific subcellular localizations. Evidence for subpopulations of organelles has been obtained for synaptic vesicles^2^, mitochondria ^3^, and insulin secretory vesicles^4^, among others. The different subpopulations of organelles may represent differences in age, maturation, or even function because of variations in biochemical composition. To better characterize cell function, we must identify the number of organelle subpopulations that exist and their distinct functional states. Understanding the dynamics of organelle composition and identifying functional subpopulations is a fundamental aspect of cell biology that is limited by lack of appropriate experimental methods. To address this gap in technology, we developed a new approach for organelle isolation. Ultimately, this approach will be useful for follow up biochemical analyses such as proteomics, functional assays, and imaging.

Conventional separation methods have proved powerful for numerous biological analyses. This process traditionally consists of iterative centrifugation steps^5^ to isolate populations of targeted organelles for follow up analysis with mass spectrometric or other omics-scale assays^6^. Gradient columns have successfully isolated clathrin-coated vesicles^7^, lipid droplets^8^, mitochondria, and endoplasmic reticulum (ER) populations^9^, from other organelles, based on differences in organelle densities. However, subpopulations of a given organelle with similar buoyant densities are often hard to differentiate and separate in these columns. Immunoisolation of organelles, although yielding pure populations, is limited by the need for specific organelle markers^10^. Electrophoretic approaches, such as Free-Flow Electrophoresis (FFE), have been adopted for use in micro-fluidic separation of organelles^11^, enabling downstream analyses of the enriched fractions in the absence of major contaminants; however, such electrophoretic separations rely on charge differences of the components. Although powerful, each of these traditional isolation approaches lack sufficient sensitivity and robustness to isolate subpopulations of an organelle, except for smooth versus rough ER^12^, for biochemical characterizations such as proteomic analysis or imaging. Thus, an alternative approach to organelle isolation will be beneficial to define organelle subpopulations more accurately.

Direct current insulator-based dielectrophoresis (DC-iDEP) provides a valuable tool for separating subpopulations of bioparticles with high resolution including viruses, bacteria, organelles, and proteins^13-16^. This approach offers a wide dynamic range, as it has been used for separations ranging from neural progenitors and stem cells^17^ to resistant versus susceptible strains of cellular pathogens^15,18^. In DC-iDEP, subtle biophysical differences can be distinguished by the differences in dielectrophoretic (DEP) and electrokinetic (EK) forces that result from a rich set of distinguishing factors associated with the radius, zeta potential, permittivity, interfacial polarizability and conductivity of the particles, to name but a few^18-22^. DC-iDEP is well-suited for assaying subpopulations of organelles, where differentiating factors are not known, because it is unbiased compared to approaches that rely on defined biomarkers. DC-iDEP can be used as a discovery-based approach to interrogate a broader spectrum of organelle subpopulations^17^ because the bioparticle separations occur quickly and the required sample volume is small^14,23,24^. Moreover, DC-iDEP is extremely sensitive to subtle biophysical differences of bioparticle or organelle identities and has been shown to be reproducible^15,17^. Here, we demonstrate the power of DC-iDEP in organelle isolation, by using it to investigate subtle differences in the subpopulations of insulin vesicles in pancreatic β-cells.

Pancreatic β-cells are responsible for secreting insulin in a tightly regulated process that is key to maintaining glucose homeostasis. Insulin vesicles undergo a complex functional maturation process that is required for proper secretion of insulin and this process is dysregulated in disease state such as diabetes^4^. Immature insulin vesicles act as a sorting compartment^25,26^ and mature into two distinct pools of functional vesicles within the cell: the readily releasable pool and the reserve pool^27-30^ (Fig. 1A). This apparent heterogeneity among subpopulations of insulin vesicles is thought to arise in part from the maturation process of insulin vesicles^31,32^ as well as their age, mobility, localization and modifications of insulin vesicle membrane proteins within the cell^4,33-35^. However, the associated biochemical constituents of insulin vesicles have remained elusive in the absence of sensitive isolation approaches. The importance of insulin vesicles in glucose homeostasis has led several groups to attempt to isolate and characterize the insulin vesicles^36,37^. While these studies and follow-up proteomics analysis of the isolated vesicles^39-42^ have provided insights into the biochemistry of these vesicles, there’s very little overlap in protein IDs associated with these organelles from different studies (Fig. 1B). Thus, there is a need for more robust isolation methods that can reproducibly differentiate between the heterogeneous subpopulations of insulin vesicles, among other organelles, and allow for their downstream characterization.

**Figure 1.**
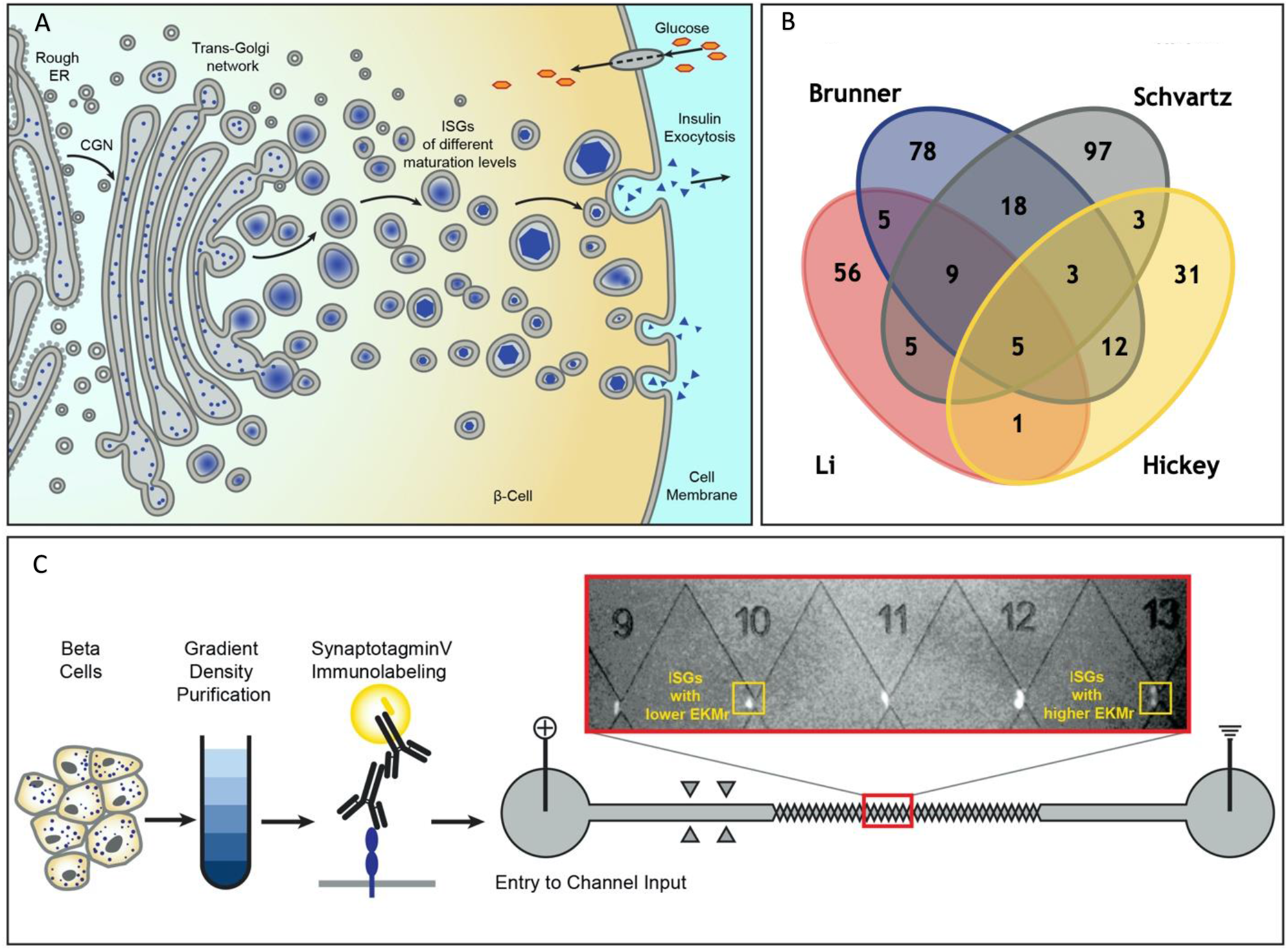
Schematic diagram for the formation of heterogenous insulin vesicles in pancreatic β-cells, graphical summation of disparate protein identifications, and processing of insulin vesicles including DC-iDEP device. A, insulin vesicle formation and maturation in a pancreatic β cell. Newly synthesized insulin is packed inside secretory vesicles which mature to store crystalline insulin in vesicles until secretion is stimulated through different signaling pathways. B, four published insulin vesicle proteomics studies ^39-42^ aimed to identify the proteome of the heterogenous populations of secretory vesicles in β-cells with only 5 proteins identified consistently. C, separation of insulin vesicles using a DC-iDEP device. Differential and density gradient centrifugation were used to enrich each sample for insulin vesicle populations. Samples were then immunolabeled and introduced into DC-iDEP device for high resolution separation. Fluorescently labeled particles trapped near various gates in the channel are biophysically different subpopulations with varied EKMr (see text) values. The gates were constricted by increasing sizes of paired triangles, forming channel widths of 73 µm to 25 µm from inlet to the outlet. The different gates created 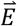 and 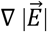 distributions for EKMr values.

In this study, a new scanning voltage DC-iDEP separation strategy has been applied to immunolabeled insulin vesicles of the INS-1E β-cells (rat insulinoma cell line^43^) and has been shown to separate the full range of insulin vesicle subpopulations with improved resolution within multiple ranges of biophysical parameters (Fig. 1C). We demonstrate that glucose treatment, which has been shown to influence the dynamics of this organelle within the β-cell^44^, affects the biophysical characteristics associated with the vesicle subpopulations captured within our DC-iDEP device. This method allows discovery of subpopulations with distinct biophysical properties amongst insulin vesicles from untreated cells (n-insulin vesicles) and 25 mM glucose-treated cells (g-insulin vesicles). Our observations are consistent with previous studies where a pronounced shift in the molecular density of the insulin vesicles was noted under the two conditions^44^. This study substantiates the sensitivity of DC-iDEP separation technique in resolving subpopulations of insulin vesicles, among other organelles, and opens the avenue for numerous studies of the biochemical constituents where complex and heterogeneous populations of organelles are of interest.

## METHODS

No statistical methods were used to predetermine sample size. The experiments were not randomized, and investigators were not blinded to allocation during experiments and outcome assessment.

### Cell culture

INS-1E cells (Addex Bio C0018009; RRID (Research Resource Identifiers) accession number: CVCL_0351) were cultured according to the supplier’s protocol. Briefly, cells were seeded in RPMI 1640 media (modified to contain 2 mM L-glutamine; 10 mM HEPES pH 7.2, 1 mM sodium pyruvate, 2 g/L glucose, and 1.5 g/L sodium bicarbonate, 50 µM 2-mercaptoethanol, 100 U/mL penicillin, and 100 µg/mL streptomycin; sterile filtered through 0.22 µm filter) supplemented with 10% fetal bovine serum (FBS) and grown to 80% confluency. Cells were plated at a density of 10^5^ cells/cm^2^ in a 24 well plate for the glucose sensitivity assay and incubated at 37°C with 5% CO_2_ in growth media for 4-5 days. Cells were pretreated at 60-80% confluency with Krebs-Ringer bicarbonate HEPES (KRBH) buffer (135 mM NaCl, 3.6 mM KCl, 5 mM NaHCO_3_, 0.5 mM NaH_2_PO_4_, 0.5 mM MgCl_2_, 1.5 mM CaCl_2_, 10 mM HEPES, pH 7.4, and 0.1% Bovine Serum Albumin (BSA); made fresh within 7 days of use) free of glucose and incubated at 37°C with 5% CO_2_ for 30 min. They were then stimulated for 30 min at 37°C using KRBH buffers containing 1.1, 5.6, 8.4, 11.1, 16.7, and 25 mM glucose, in the presence of house-made protease inhibitor (PI) cocktail (0.5 M AEBSF, 1 mM E-64, 1.13 mM leupeptin, and 151.36 μM aprotinin). KRBH buffer was removed and saved from cells for downstream analysis. Insulin secretion in response to increasing concentrations of glucose was confirmed in an Enzyme Linked Immuno-Sorbent Assay (ELISA) (Mercodia 10-1250-01) following the manufacturer’s manual (Fig. S1). For each biological replica, cells were plated at a density of 4×10^4^ cells/cm^2^ in a 5-layer cell chamber (VWR 76045-402) to yield enough material for completing the experiment. All cell stacks were at least 95% viable after the harvest with 0.05% trypsin. For glucose treatment, near 80% confluent cells were gently rinsed with dialyzed phosphate-buffered saline (dPBS) twice and starved in a KRBH buffer with no glucose for 30 min, followed by a 30 min stimulation of insulin release by KRBH buffer supplemented with 25 mM glucose. Cells were then harvested by mild trypsinization.

### Colocalization of synaptotagmin V and insulin vesicles

Cells were grown on Nunc Lab-Tek II Chamber Slide, precoated with poly-L-lysine. Cells were fixed with 4% ice-cold PFA for 10 min, and then stained with antibody cocktail (Rabbit anti-insulin antibody (Abcam ab181547), 1:1000; Mouse anti-synaptotagmin V (BD Biosciences, 612284), 1:1000) in 0.5% BSA, 0.2% saponin, 1% FBS of PBS buffer for 1 h at room temperature (RT). After three washes with PBST for 10 min, cells were incubated for 40 min with secondary antibody cocktail (Goat anti-rabbit IgG H&L, Alexa 488 (Abcam ab150077) 1:1000; Goat anti-Mouse IgG (H+L) Cross-Adsorbed Secondary Antibody, Alexa Fluor 647 (Invitrogen A-21235), 1:1000). Cells were then mounted in ProLong™ Glass Antifade Mountant with NucBlue™ Stain (Thermo Fisher Scientific P36981) and cured for 24 h before imaging.

### Microscopy imaging

A Leica SP8 FALCON with DIVE laser scanning microscope was used for confocal imaging. Image acquisition was performed using 63x/1.4NA oil immersion objective, using 1024×1024 format at 400 Hz, with 0.03 μm pixel size, and pinhole set to 1 AU. The signal was collected with 405 nm, 499 nm, and 653 nm excitations and 410-494 nm, 504–621 nm, 658-776 nm emissions for nuclei, insulin, and synaptotagmin V by HyD detector. The images were deconvoluted by LAS X, lightening module. The refraction index was set to 1.52 for processing.

### Insulin secretory vesicle enrichment

Following trypsinization, all the steps were performed at 4°C. Cells were gently washed twice with PBS, followed by Dounce-homogenization of the cells with 20 strokes in homogenization buffer (HB) (0.3 M Sucrose, 10 mM MES, 1 mM EGTA, 1 mM MgSO_4_, pH 6.3) supplemented with house-made PI cocktail (0.5 M AEBSF, 1 mM E-64, 1.13 mM leupeptin, and 151.36 μM aprotinin). Cell debris was collected by centrifugation at 600x g for 10 mins and re-homogenized as described above to lyse the remaining intact cells, followed by a second spin at 600x g. Supernatants were pooled and centrifuged at 5,400x g for 15 min to remove mitochondria, ER, and other subcellular compartments of similar density. The pellet was discarded, and the supernatant was centrifuged at 35,000x g for 30 min to sediment insulin vesicles, among other contaminants, yielding up to ∼5 µg of dry material per every million cells. This pellet was resuspended in ∼450 μL HB and loaded on a density gradient column formed by layering decreasing densities of OptiPrep density media (Sigma-Aldrich D1556) and HB in 0.9 mL fractions of 40, 35, 30, 25, and 20 percent OptiPrep in an open-top thin-wall polypropylene tube (Beckman 326819). The density column was then spun in a SW55i Beckman rotor of an ultracentrifuge at 160,000x g for 8 h to fractionate the insulin vesicle-containing population. Insulin vesicle subpopulations were isolated in fractions of 400 µL, and ELISA and western blotting (WB) were used to identify the fractions most enriched in insulin. A similar dilution was applied to all the fractions. The manufacturer’s manual was followed for ELISA (Mercodia 10-1250-01), (Fig. S2A). For WB, fraction samples were mixed with 4X NuPAGE™ LDS sample buffer (Invitrogen NP0007), loaded on a 15-well NuPAGE™ 4 to 12% bis-tris gel (Invitrogen NP0323PK2) and run in a mini-gel tank (Life Technologies A25977) at 200V for 30 min (Bio-Rad 1645050). Protein was then transferred to a PVDF membrane using iBlot™ 2 Transfer Stacks (Invitrogen IB24002) in iBlot 2 dry blotting device (Invitrogen IB21001). The membrane was blocked using 5% BSA, then cut according to marker protein size and incubated with antibodies against marker proteins for mitochondria (Cytochrome c Antibody; Novus Biologicals NB100-56503), ER (SEC61B Polyclonal Antibody; Life Technologies PA3015), and insulin vesicles (synaptotagmin V; Thermo Fisher Scientific PA5-44987; RRID: AB_2610517) at RT for 2-5 h. Membranes were then washed with 0.1% Tween supplemented PBS (PBST) twice and incubated with the secondary antibody ((anti-rabbit IgG), (anti-mouse IgG)) at RT for 1 h. Membranes were washed with PBST 2-3 times and bands were visualized upon addition of SigmaFast BCIP^®^/NBT tablets (Sigma B5655) (Fig. S2B). Fractions containing the highest insulin levels and high concentration of the vesicle marker synaptotagmin V were regarded as insulin vesicle samples and were further tested for dynamic light scattering (DLS) using Wyatt Technology’s Mobius to confirm the size distribution of particles corresponding to the insulin vesicle diameter, reportedly 200-500 nm. Data was analyzed in DYNAMICS and manually corrected against the control (HB) (Fig. S2C). Insulin vesicle fractions confirmed to have the expected insulin vesicle diameter by DLS were then pooled together and spun down at 35,000x g for 15 min to sediment the insulin vesicles.

### Immunolabeling of insulin vesicles

The pellet was resuspended in HB and incubated with anti-synaptotagmin V (Thermo Fisher Scientific PA5-44987; RRID: AB_2610517) to tag an insulin vesicle marker, which served as a primary antibody that was further stained with Alexa 568-conjugated secondary antibody (Invitrogen A-11011; RRID: AB_143157) (Fig. 1). Insulin vesicles were washed with HB three times to remove the excess antibody and were finally resuspended in a low conductivity buffer (LCHB; 0.3 M sucrose, 5 mM MES, pH 6.3), which is compatible with dielectrophoresis studies.

### Device fabrication

The design and fabrication methods of the separation device were described in prior publications^45^. The device contains a 27-gate sawtooth channel with a depth of approximately 20 μm and a length of 3.5 cm from inlet to outlet (Fig. 2C). The distance between two paired triangle tips (gates) decreases from 73 μm to 25 μm in the channel. The gate size decreases approximately 5 µm after every 3 repeats. Direct current was applied to the device between the inlet and outlet. The potentials were between 0–1800V for testing.

**Figure 2.**
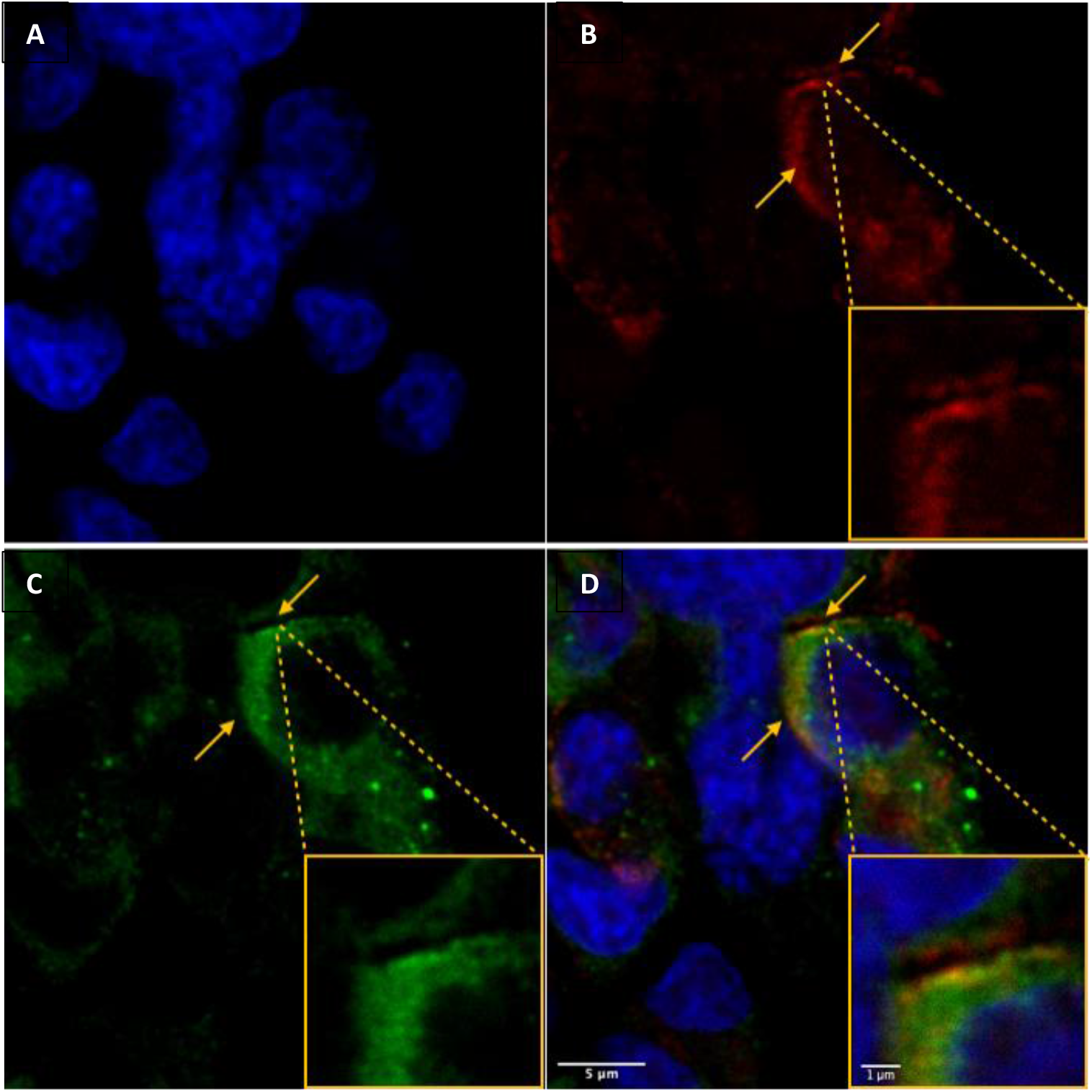
Colocalization of insulin vesicle marker used in this study (synaptotagmin V) with insulin. A, nuclei of INS-1E β-cells stained with NucBlue™. B, synaptotagmin V labeled with mouse anti-synaptotagmin V and goat anti-mouse IgG (H+L), Alexa 647. C, hormone insulin labeled with rabbit anti-insulin and goat anti-rabbit IgG (H+L), Alexa 488. D, localization of synaptotagmin V to insulin vesicles as apparent from the merged intensities of panels B and C. Microscopy was performed with a Leica microscope using a 63x/1.4NA oil immersion objective on cells mounted in ProLong™ Glass Antifade Mountant.

The microfluidic devices were fabricated by standard soft lithographic technique as described previously^45^. The design of the channel was created by AutoCAD (Autodesk, Inc., San Rafael, CA) and was used for fabricating a photomask. The channel was created by exposing AZ P4620 positive photoresist (AZ Electronic Materials, Branchburg, NJ) on Si wafer CEM388SS (Shin-Etsu MicroSi, Inc., Phoenix, AZ) by contact lithography. Extra materials were removed from the Si wafer. A weight of 22 g of polydimethylsiloxane (PDMS, Sylgard 184, Dow/Corning, Midland, MI) was used to fabricate four channels simultaneously. The PDMS mixture was placed on the Si wafer template and left to stand for 30 min to allow bubbles to dissipate, and then it was baked for 1 h at a temperature of 80°C. Holes 2.5 mm in diameter were punched for inlet and outlet reservoirs. Each channel was capped with a glass microscope slide to fabricate the enclosed channels after cleaning and activation by plasma cleaner (Harrick Plasma, Ithaca, NY, USA) with a voltage of 50 kV.

### Electric field simulations

Finite element modeling (COMSOL, Inc., Burlington, MA) of the distribution of the electric field in the microchannel was performed as previously detailed^45^. The *AC/DC module* was used to interrogate the 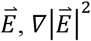, and 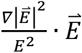 in an accurately scaled 2D model of the microchannel.

### DC-iDEP

The separation channel was treated with 5% (w/v) BSA for 15 min followed by a wash with LCHB. A volume of 15 μL of the insulin vesicle sample was introduced to the device from the inlet. This sample fraction contains particles ∼75% of which have radii characteristic of insulin vesicle as apparent from DLS experiments (Fig. 2SC). The volume in each reservoir was maintained with LCHB to prevent pressure induced flow. Direct current was applied at 600V, 900V, 1200V, 1500V, and 1800V between the inlet and outlet and particles were driven through the channel by the EK force experienced.

### Imaging

Images and recordings were acquired using an Olympus IX70 inverted microscope with 4X, N.A. 0.16, and 20X, N.A. 0.40, objectives. The 20X objective was used to inspect the channel to confirm that the device was properly formed. The 4X objective was used in recording the intensity of the insulin vesicle signal shown in Figure 1. A mercury short arc lamp (H30102 w/2, OSRAM) and a triple band pass cube (Olympus, Center Valley, PA) were used for sample illumination and detection (Excitation:400/15-495/15-570/25; Dichroic:410-510-590; Emission:460/20-530/30-625/50). Fluorescent intensities of immunolabeled insulin vesicles were recorded using the 4X objective by a LightWise Allegro camera (LW-AL-CMV12000, USB3, 0059-0737-B, Imaging Solutions Group) after the voltage had been applied 90 s. Images were recorded from 3-4 biological replicates at each gate (27 total gates) for any given voltage. Images were further processed in ImageJ (NIH, freeware). The intensity at each gate was recorded along with the intensity of a nearby open area of the channel (image intensity background), which was subtracted from the intensity at each gate to adjust for any variation in illumination intensity. The data for each applied voltage value was normalized to the highest intensity within that data set.

### Theory

The forces exerted on bioparticles in the microfluidic device in the presence of direct current include dielectrophoretic (DEP) and electrokinetic (EK) forces. Separation of subpopulations is achieved based on the different magnitude of the forces each bioparticle experiences, related to the properties of the particles, including their radius, conductivity, and zeta potential.

The EK mobility, *µ*_*EK*_, and velocity, 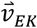, are described as:

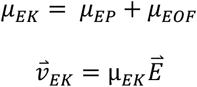

where *µ*_*EP*_ is the electrophoretic (EP) mobility and *µ*_*EOF*_ is the electro-osmotic flow (EOF) mobility. The DEP mobility, *µ*_*DEP*_ and velocity, 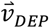 can be expressed as:

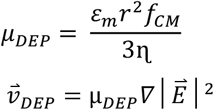

where *r* is the radius of the particle, *f*_*CM*_ is the Clausius-Mossotti factor, *ε*_*m*_ is the dielectric constant of the solution, and *η* is the viscosity.

A combination of biophysical properties of the particles, such as insulin vesicles, determines the location where they will be captured in a microfluidic device. Capture occurs when the EK velocity of the particle is equal to that of DEP. The condition is:

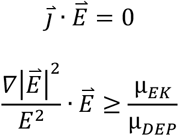

where 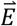 is the electric field intensity, 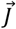 is the particle flux, and 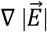 is the gradient of the electric field. The ratio of EK to DEP mobilities 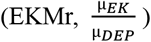 can be used to characterize the biophysical properties of different subpopulations^17^. The EKMr of an insulin vesicle is larger than 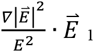 of the gates which it has passed through and is smaller/equal to 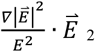 of the gates where it is captured. In this way, insulin vesicles are separated in the microfluidic channel, thus measuring the EKMr values for the insulin vesicles.

## RESULTS

### Analysis of enriched insulin vesicle samples post fractionation

The INS-1E cells used in these studies were capable of insulin secretion in response to increasing concentrations of glucose (Fig. S1). Isolated membrane fractions from differential and density gradient centrifugations were analyzed through ELISA, western blotting (WB) and dynamic light scattering (DLS). ELISAs identified that lower fractions of the density column (9-12) contained the highest insulin content (Fig. S2A), the same fractions shown to be enriched in the insulin vesicle marker, synaptotagmin V, by WB (Fig. S2B). DLS indicated that these fractions contained particles of 150-200 nm radius, which corresponds to the known range of insulin vesicles radii^46,47^. Although the WB revealed that the final enriched vesicle sample included some contaminants from unwanted organelles, such as the ER (Fig. S2B), this will not confound the analysis of insulin vesicles in the iDEP device as the fluorescent labeling was targeted only at insulin vesicles.

### Introduction of DC-iDEP as a discovery and quantification tool for insulin vesicle subpopulations

Enriched vesicle samples from INS-1E cells were subjected to analysis using DC-iDEP after labeling with insulin vesicle marker synaptotagmin V^39,41^, which was confirmed to co-localize with insulin in fluorescence microscopy imaging (Fig. 2). The separation system was operated in a discovery or scanning mode by first applying a high voltage (2100V, empirically determined, Fig. 3) such that all particles were prevented from entering the first gate, because DEP forces exceed EK forces. The voltage was then lowered incrementally (−300V for each step) allowing various subpopulations to enter the separation zone and be sorted along the device according to their specific EKMr values (Fig. 3). A bolus forms at a gate, corresponding to an EKMr value that is a result of a balance between DEP and EK forces on each particle and reflects a complex set of biological, chemical, and biophysical properties of the vesicles^15,48-50^. The fluorescence intensity was captured for each gate over a full range of voltages (1800V to 600V), such that the largest EKMr values are probed with the higher applied voltage (Fig. 4).

**Figure 3.**
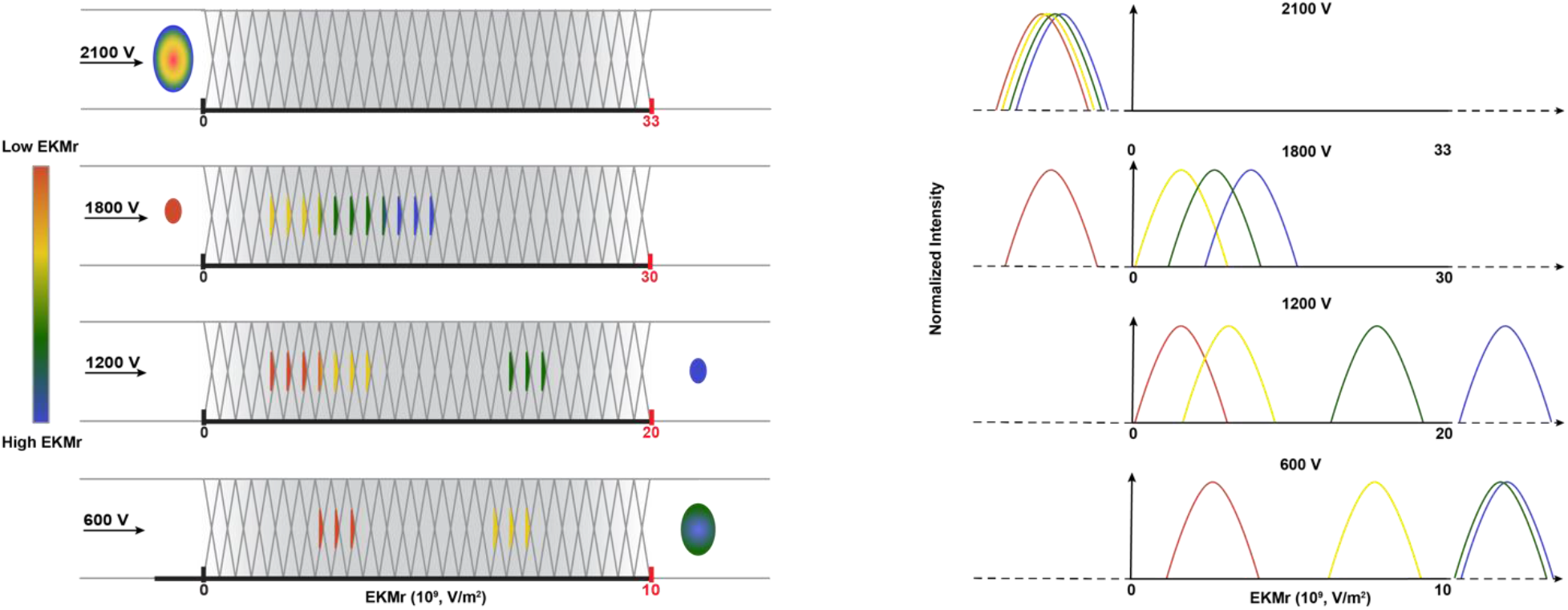
Schematic diagram of DC-iDEP system operated in a discovery or scanning mode. A, as a function of the design of the sawtooth channel with different gate sizes along the channel, the applied voltage defines DEP and EK forces at each gate, and capture occurs when the EK force of the particle is equal to or smaller than the DEP force. At high voltages, only particles with high EKMr values can enter the channel; the highest applied voltage of 2100V prevents all particles of the sample from entering the inlet of the device due to the induced dielectrophoretic forces (no fluorescent signal detected anywhere along the channel). Sequentially lower voltages allow the various subpopulations to enter and be separated throughout the channel. When a subpopulation’s EKMr value surpasses the channel’s DEP force limit, it travels freely and leaves the channel at the outlet. B, fluorescent intensities of the captured particles are recorded along the channel at each voltage. Tracking these intensities allows the discovery and quantification of unknown subpopulations according to their biophysical properties.

**Figure 4.**
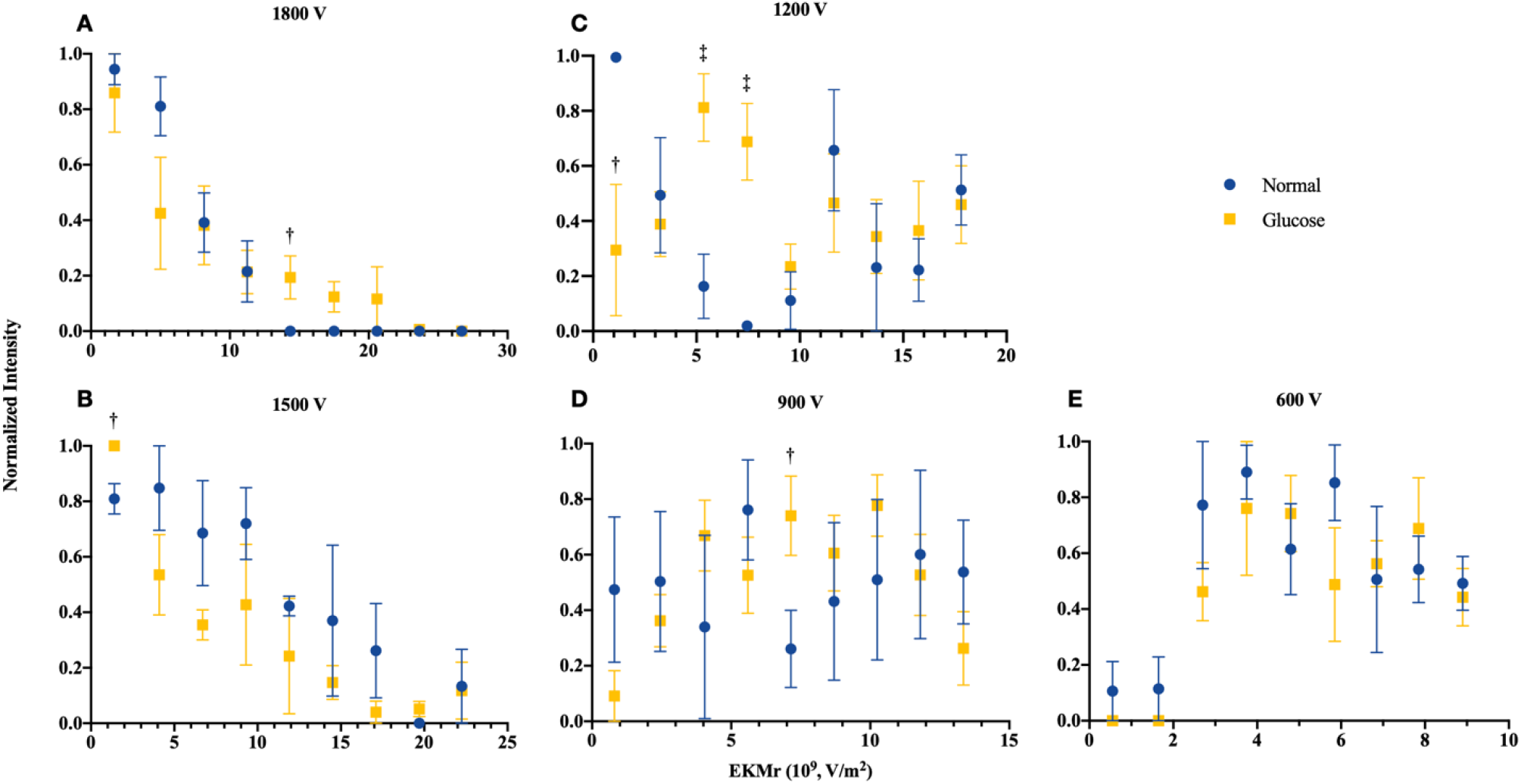
Comparison of distributions for the biophysical properties as reflected in EKMr values of n-insulin vesicles (blue circles) and g-insulin vesicles (yellow squares) with varied voltages applied. Fluorescent intensities of captured n- and g-insulin vesicles with different EKMr values were recorded at each gate. Each data point reflects fluorescent intensities recorded at 3 subsequent gates with the same EKMr values, averaged out over biologically replicated experiments and normalized over all signals recorded at a given voltage. Full profile of the sample’s biophysical distribution was recorded at 1800V. Insulin vesicle subpopulations were separated at subsequent applied voltages of 1500, 1200, 900, and 600V. Values are mean ± SEM (n = 3 for n-vesicles or 4 for g-vesicles for biologically independent experiments). The two sample Cramer-von Mises Criterion shows that there is a larger than 99.9% probability that the n-insulin vesicles and g-insulin vesicles distribution differs at all voltages. Data are unity-based normalized and analyzed using student t-test to determine significant difference between data points of the two conditions. The t-test shows the n- and g-vesicles from the 3.25-7.45 V/m^2^ EKMr range (C) differ significantly at 95% confidence level (‡P<0.05). This test also shows that at the 90% confidence level (†P<0.10), the n- and g-vesicle data points differ at the EKMr ranges of 0-1.40×10^9^ V/m^2^ (B-C), 5.60-7.15×10^9^ V/m^2^ (D), and 11.25-14.35×10^9^ V/m^2^ (A).

### Biophysical subpopulations of vesicles from untreated β-cells

The distribution of fluorescently labeled n-insulin vesicles captured at each gate formed a characteristic arc (Fig. 1C), indicating a well-operating and consistent system. Higher voltages provide a broader dynamic range of EKMr values for capturing a wider range of particles, while lower voltages provide detailed distribution of particles based on their associated EKMr values. At an applied voltage of 1800V, particles were sensed at EKMr values below 1.5 ×10^10^ V/m^2^ in patterns of overlapping subpopulations (Fig. 4A, blue circles). These overlapping features begin to spread out with an applied voltage of 1500V. At incrementally lower settings of applied voltages (1200, 900, and 600V), distinctive patterns become identifiable (Fig. 4B-E). Notable and discernible features of bioparticle distribution are apparent around 1.2×10^10^ V/m^2^ and 1.8×10^10^ V/m^2^ at an applied voltage of 1200V (Fig. 4C). Lowering the voltage to 900V, and redistribution of bioparticles based on adapted properties of the channel, reveals a similar but attenuated feature of the distribution around 1.2×10^10^ V/m^2^ (Fig. 4D), whereas particles with EKMr values greater than 1.5×10^10^ V/m^2^ leave the channel at this voltage. At this voltage, redistribution of particles, previously retained in overlapping patterns at 1200V, forms a distinct peak around 5-6×10^9^ V/m^2^ (Fig. 4D). Further lowering the voltage to 600V shows similar patterns of particle distribution around 5-6×10^9^ V/m^2^, as well as distinctive patterns around 3-4×10^9^ V/m^2^ (Fig. 4E), while leaving out populations with EKMr values higher than 1.0×10^10^ V/m^2^.

### Biophysical subpopulations of vesicles from glucose stimulated β-cells

Insulin vesicles obtained from 25 mM glucose-treated INS-1E cells (g-insulin vesicles) were studied with the same method used for the untreated cells (Fig. 4, yellow squares). Consistent with the vesicles from untreated cells, patterns of primarily overlapping subpopulations were detectable at a voltage of 1800V. Distribution of particles were observed at values up to 2.3×10^10^ V/m^2^ (compared to a maximum value of 1.5×10^10^ V/m^2^ for the untreated populations) (Fig. 4A), which suggests that these vesicles have a broader range of properties than the population from the untreated cells. The first evidence of a distinct distribution feature was captured around 7-8×10^9^ V/m^2^ when the voltage was lowered to 1500V (Fig. 4B). Decreasing the voltage to 1200V and subsequent particle redistribution in the channel revealed a distribution pattern with discernible features around 7-8×10^9^ V/m^2^, like those observed at 1500V, as well as a distinct peak around 1.1×10^10^ V/m^2^ (Fig. 4C). Further lowering the voltage to 900V resulted in a unique distribution pattern with discernible features around 1.1×10^10^ V/m^2^ (Fig. 4D), similar to 1200V, and 8×10^9^ V/m^2^ (Fig. 4D), previously observed at 1500V and 1200V (Fig. 4B-C). Another feature of this distribution pattern was a peak around 4×10^9^ V/m^2^ (Fig. 4D). Ultimately, decreasing the voltage to 600V revealed a distribution pattern with features around 4×10^9^ V/m^2^ and 8×10^9^ V/m^2^ (Fig. 4E), like those observed at higher voltages, leaving out patterns that were observed at EKMr values higher than 1.0×10^10^ V/m^2^. This distinct distribution pattern, although consistent with distribution patterns at higher voltages, featured stretched-out peaks, consistent with smaller and more refined range of EKMr values assigned throughout the channel.

### Behavior comparison of insulin vesicles and glucose stimulated insulin vesicles

Visual inspection of the collected data (Fig. 4) indicated clear differences in the patterns of the insulin vesicle distributions between the vesicle samples from treated and untreated cells. The n-insulin vesicles were discernible at ∼1.2×10^10^ V/m^2^, ∼5-6×10^9^ V/m^2^, and ∼3-4×10^9^ V/m^2^ (weak at ∼1.8×10^10^ V/m^2^), whereas the g-insulin vesicles were observed at ∼1.1×10^10^ V/m^2^, ∼8×10^9^ V/m^2^, and ∼4×10^9^ V/m^2^ (weak at ∼2.1×10^10^ V/m^2^). Beyond visual inspection, two statistical methods were used to quantify and more deeply explore both the global and local differences: 1) the likelihood of the two multi-component samples being different in composition, and 2) direct comparisons of paired data points between the n-insulin vesicles and g-insulin vesicles data sets. To aid in deciding whether two complex data sets differ, the two-sample Cramer–Von Mises criterion was used^17,51^. In essence, this statistically examines the whole data set for the largest differentiation without necessarily identifying the specifics. The criterion is based on the following equation (where the resulting values of *T* can be correlated to a probability of the null result):

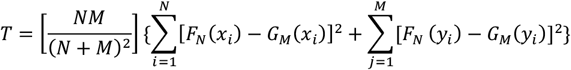

where *F*_*N*_ (*x*_*i*_) and *G*_*M*_ (*x*_*i*_) are the empirical distributions of the n- and g-insulin vesicles samples, the variables *x*_*i*_ and *y*_*i*_ are the observed values in each sample, and *N* and *M* represent the number of data points used to compare n- and g-insulin vesicles samples. An individual assessment was made for each applied voltage and indicated significant differences, consistent with the visual inspection. The calculated *T* values were 4.8×10^3^ at 600V, 4.7×10^3^ at 900V, 1.3×10^4^ at 1200V, 4.7×10^3^ at 1500V and 1.0×10^4^ at 1800V. These values indicate that at all voltages, there is a larger than 99.9% confidence level that the null hypothesis that the two samples come from the same distribution can be rejected. This confidence level is even closer to 100% at 1200V and 1800V than other voltages. Further, a student’s t-test^52^ for paired data points of the intensity at each gate confirmed a statistical difference between n- and g-vesicles from the EKMr ranges of 3.25-7.45 V/m^2^ at 1200V (Fig. 4C), 0-1.40×10^9^ V/m^2^ at 1500V and 1200V (Fig. 4B-C), 5.60-7.15×10^9^ V/m^2^ at 900V (Fig. 4D), and 11.25-14.35×10^9^ V/m^2^ at 1800V (Fig. 4A).

## DISCUSSION

Using the DC-iDEP separation method, we identified distinct distribution patterns of insulin vesicles with varying biophysical properties which we interpret as indicative of vesicle subpopulations. Furthermore, we observed different distribution patterns for insulin vesicles isolated from glucose-stimulated versus unstimulated cells, which suggests glucose stimulation alters the insulin vesicle subpopulations. This is consistent with findings from single cell analysis using cryo-electron tomography and soft X-ray tomography, which revealed heterogeneities in the composition of insulin vesicles^53^ and their molecular densities^44^ depending on the drug treatment and location of insulin vesicles within the cell. Other studies have also observed enrichment of certain subpopulations of vesicles in response to glucose stimulation^54^. Distinct subpopulations that were observed in both n- and g-insulin vesicles across a wide range of EKMr values indicate that both n- and g-insulin vesicles are made up of complex and heterogenous subpopulations and are statistically different in their overall distributions pattern as well as in several specific EKMr values. Our approach addresses an important need to identify and separate distinct subpopulations of insulin vesicles which can allow for investigating their apparent heterogeneity and further characterizing them.

The pattern of EKMr values is influenced by individual insulin vesicles. The distribution of this pattern reflects differences in physical properties of the vesicles in response to exposure of the cells to glucose. The specific changes to the individual insulin vesicles which result in varied EKMr measurements can be associated with any alteration of the biochemical makeup of the bioparticle. There is an ongoing evolution of the theoretical underpinnings of electric field gradient techniques, where past physical descriptors were limited to conductivity and permittivity of the particle.^20^ It is becoming better understood that the overall structure and subtle details of the bioparticle, including the particle-solvent cross polarizations, will influence these forces^20-22,55,56^. This new view of the forces imparted on the insulin vesicles aligns with a rather simple proposition that the makeup of the particle has changed: something has been added, subtracted, or altered which changed the force on the particle in a measurable and quantifiable way. This is consistent with findings that glucose can affect the maturation of existing vesicles, increasing their molecular density or the concentration of biomolecules within the vesicle lumen^44^. Additionally, the subpopulations we observe could have been altered by a change in surface protein expression or enrichment of unsaturated lipids^57^. This is also consistent with findings that glucose enhances vesicle-mitochondria association which is hypothesized to contribute to insulin vesicle maturation^44^. Identical particles always have the same EKMr in the same manner that identical proteins always have the same molecular weight^58^.

There are some subtleties in the presented data which accentuate features, and some limitations, of the DC-iDEP separations and the imaging system. The data for 1800V, for instance, shows no discernable local maxima above 1.5×10^10^ V/m^2^ for the n-vesicles, despite having identifiable populations with EKMr values higher than 1.5×10^10^ V/m^2^ at lower voltages. Another feature that is quite apparent is a lack of distinct ‘peaks’ or identifiable patterns appearing at consistent EKMr values within the data sets from differing applied voltage values. While the applied voltage does not affect the properties of bioparticles, it defines the forces that oppose bioparticle movements across the channel at different gates. Accordingly, in an overlapping pattern of subpopulations, those with higher EKMr values overcome the weaker opposing forces at a given gate once the voltage is lowered.^16,59^ Hence, the signal which was previously averaged with signals from the other overlapping subpopulations is now discernible at a higher EKMr value. Considering the high resolution of the technique, homogeneous subpopulations of vesicles undoubtably consist of a very narrow range of EKMr values. Each data point shown in Figure 4 represents a homogenous subpopulation captured at the gate. While this feature is unsatisfying to classic separation scientists, it still allows for quantitative comparison of paired samples, as has been shown here. To further optimize the separation, one could modify the channels including the gate size and periodicity, as well as scan even more refined ranges of voltages to induce varying spectra of EKMr values along the device ^14,15^. Additionally, developing the capability to port the collected individual boluses will enable downstream analyses such as mass spectrometry or electron microscopy.

In essence, the current work introduces DC-iDEP in a scanning mode as a powerful tool to interrogate complex organelles subpopulations and study their distinctive distribution patterns under different treatment conditions with the goal of organelle subpopulation discovery and quantification. Further, the putatively subtle differences in subpopulations between stimulated and unstimulated β-cells are quantified more extensively than previously possible. Finally, this method provides a mechanism for the isolation and concentration of fractions which show the largest difference between the two population patterns for further bioanalysis (imaging, proteomics, lipidomics, etc.) that otherwise would not be possible given the low-abundance components of these subpopulations. This approach can be broadly applied to any cell type and organelle beyond the scope of our model system of insulin vesicles and β-cells.

## Author Contribution

KLW, SEF, MAH and MB conceived the project. MB performed treatment on cells, harvested the cells, and performed insulin vesicle enrichment followed by QC with ELISA, WB, and DLS. ZW performed fluorescent microscopy. YL and MAH designed the iDEP experiments. YL performed all iDEP experiments, data reduction and presentation, and conceived the statistical analysis. MB and YL performed statistical analysis on the iDEP data. MAH conceived the Figures and MB created the Figures with input from all authors. All authors participated in data interpretation. All authors helped write and edit the manuscript, with YL and MB as leads.

## Acknowledgements

The authors would like to thank Raymond Stevens and the Bridge Institute for supporting our project and members of the Pancreatic β Cell Consortium for their inspiring discussions and feedback. We additionally thank Chris Hanson for assisting with the cell culture, Yekaterina Kadyshevskaya for helping with illustrations, Claire Cato for feedback on the manuscript, Brett Barbaro for assisting with analysis of the existing proteomics data, and USC NanoBiophysics Core Facility for facilitating the DLS experiments.

## Funding

This work was supported by the Bridge Institute at the USC Michelson Center for Convergent Bioscience (KLW) and National Institutes of Health grant 5R03AI133397-02 (MAH).

**Figure S1.**
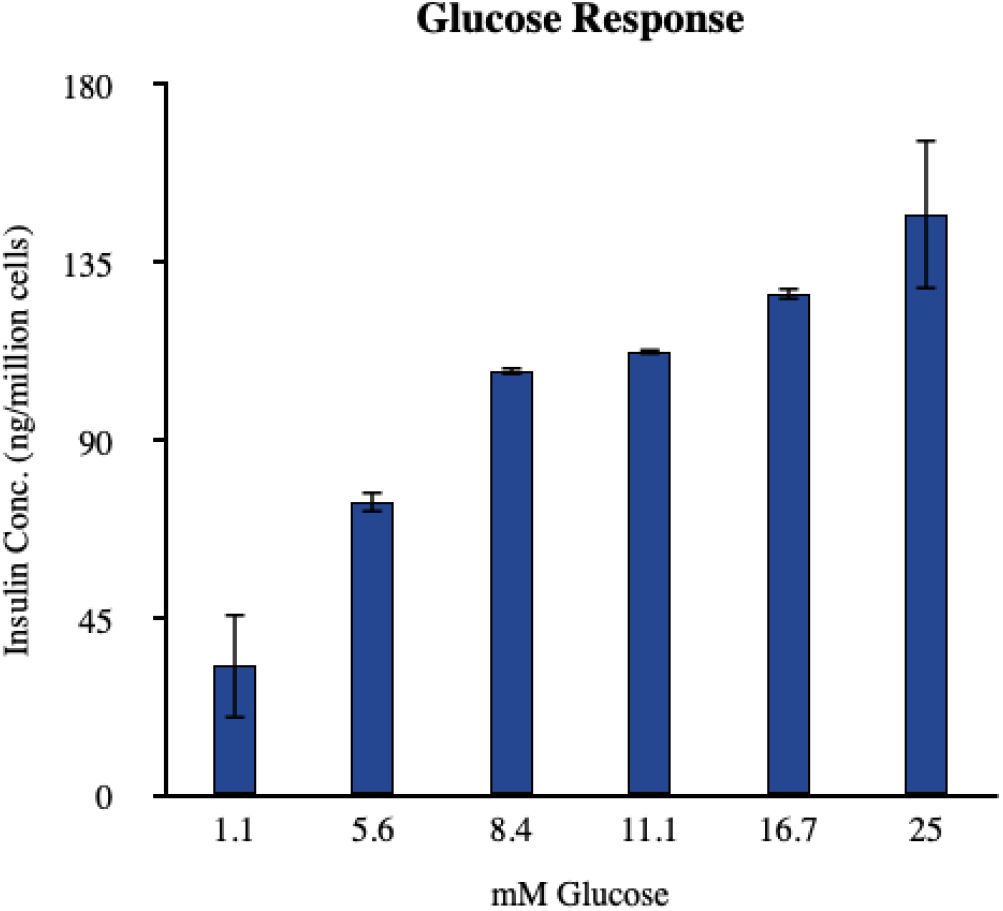
Glucose sensitivity of INS-1E β-cells was tested by stimulation at increasing concentrations of glucose and measurement of insulin secretion by ELISA.

**Figure S2.**
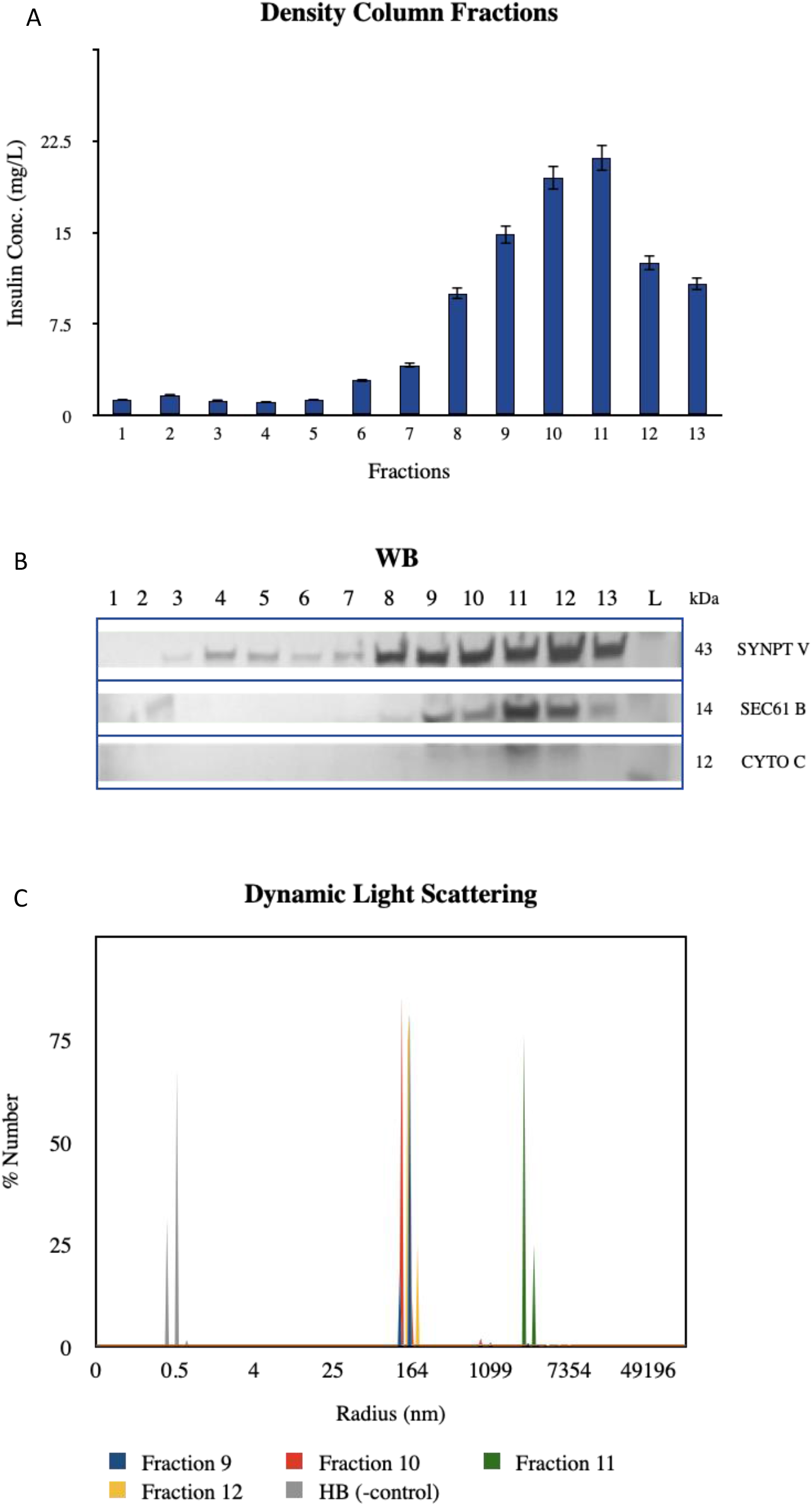
A, fractions of the density column were screened for insulin content in the ELISA assay. B, western blotting of the density column fractions revealed high concentrations of insulin vesicle marker synaptotagmin V in fractions with high insulin content as well as presence of ER and mitochondria contaminants as indicated by organelle markers SEC61 and Cytochrome C, respectively, in the same fractions. C, DLS was performed on fractions of interest to validate presence of particles of 150-200 nm in radius, corresponding to radii of insulin vesicles.

